# Adapting the novel-object recognition task to quantify how much attention rodents allocate to motivationally-salient objects

**DOI:** 10.1101/495945

**Authors:** Iasmina Hornoiu, John Gigg, Deborah Talmi

**Affiliations:** Division of Neuroscience and Experimental Psychology, University of Manchester; Department of Psychology, University of Cambridge

**Keywords:** attention, emotional arousal, memory, Novel Object Recognition task, olfactory stimuli

## Abstract

The allocation of attention can be modulated by the emotional value of a stimulus. In order to understand the biasing influence of emotion on attention allocation further, we require an animal test of how motivational salience modulates attention. In mice, female odour triggers arousal and elicits emotional responses in males. Here, we determined the extent to which objects labelled with female odour modulated the attention of C57BL/6J male mice. Seven experiments were conducted, using a modified version of the spontaneous Novel Object Recognition task. Attention was operationalised as differential exploration time of identical objects that were labelled with either female mouse odour (O+), a non-social odour, almond odour (O^a^) or not labelled with any odour (O-). In some experiments we tested trial unique (novel) objects than never carried an odour (X-). We found that when single objects were presented, as well as when two objects were presented simultaneously (so competed with each other for attention), O+ received preferential attention compared to O-. This result was independent of whether O+ was at a novel or familiar location. When compared with O^a^ at a novel location, O+ at a familiar location attracted more attention. Compared to X-, O+ received more exploration only when placed at a novel location, but attention to O+ and X- was equivalent when they were placed in a familiar location. These results suggest that C57BL/6J male mice weigh up aspects of odour, object novelty and special novelty for motivational salience, and that, in some instances, female odour elicits more attention (object exploration) compared to other olfactory stimuli and visual object novelty. The findings of this study pave the way to using motivationally-significant odours to modulate the cognitive processes that give rise to novel object recognition.

## Introduction

The Novel Object Recognition (NOR) task is one of the most widely-used paradigms in studies of rodent memory. Originally introduced in rats by Berlyne (1950) it was subsequently developed by Ennaceur and Delacour (1988) and later adapted for use in mice by Murai *et al*. (2007). The task relies on the spontaneous drive of rodents to explore novelty in the absence of any training or external reinforcers. The original NOR task comprises three phases: habituation, familiarisation and test. In the first two phases, the animal is exposed to an open-field arena (habituation) and then to the same arena in the presence of two identical objects (familiarisation). The test phase is similar to the familiarisation phase, with the exception that one of the identical objects is replaced with a novel object. Healthy adult rodents recognise the familiar object at test and explore the novel object for a longer duration than the familiar object (Ennaceur, 2010; Gaskin et al., 2010; Mumby, Glenn, Nesbitt, & Kyriazis, 2002). The recognition of a familiar object is thus expressed in the NOR in the increased exploration of the novel object.

Preferential exploration of the novel object during the test phase of the NOR task clearly indicates whether animals remember the objects that were presented in the familiarisation phase. There is less clarity on the cognitive function indexed by the duration of exploration of objects during the familiarisation phase. Piper and colleagues (Piper, Fraiman, & Meyer, 2005) suggested that the duration and frequency of object exploration during that phase could provide an index of attention to objects. It is reasonable to assume that if an animal pays more attention to objects in the familiarisation phase, namely, approaches the objects more frequently and explores them for longer, it is likely to encode these objects better into long-term memory. Consequently, these objects are likely to be recalled more reliably later, indexed through increased preference for the novel object in the test phase. Defining the extent of object exploration in the familiarisation phase as a behavioural measure of attention has face validity because it aligns with the use of this term in human research, where overt attention is often measured through the number of eye fixations (Christianson, Loftus, Hoffman, & Loftus, 1991; Kim, Vossel, & Gamer, 2013; Riggs et al., 2011). In addition, the exploration of one object in the NOR decreases the exploration of the other, reminiscent of the ‘gold standard’ of measuring human attention capacity through the degree to which the target item impairs performance on a competing task (Craik, Naveh-Benjamin, Govoni, & Anderson, 1996; Kahneman, 1973; Kern, Libkuman, Otani, & Holmes, 2005; Talmi, Schimmack, Paterson, & Moscovitch, 2007). Therefore, by measuring attention to objects during the familiarisation phase researchers may be able to distinguish between animals with attention versus memory deficits. For example, a variant of the NOR has been used in studies focusing on pathological conditions or drug abuse, which result in attentional deficits (Alkam et al., 2011; Piper, Fraiman, & Meyer, 2005). Alkam and colleagues also suggest that object exploration in the NOR could be a measure of attention especially when there is no time delay between the familiarisation and test phases. The preference to explore one object over another aligns with selective attention, and the duration of exploration, with sustained attention (Bushnell & Strupp, 2009). Nevertheless, the NOR has only seldom been used to study attention function in rodents and, therefore, its potential to illuminate human attention mechanisms has not been fully realised. Here, we present an adaptation of the NOR to investigate emotional biases in attention allocation in mice, measured through increased object exploration.

It is interesting to use the NOR for this purpose because current animal tests of attention typically employ tasks derived from theories of associative learning and conditioning, which are quite different from those used more commonly to study attention in human participants. In one model of associative learning, proposed by Mackintosh (1975), animals pay more attention to cues that are reliable predictors of a consequence than to non-predictive cues. Selective attentional bias towards good predictors allows animals to focus on relevant cues, while ignoring distractors, thereby, achieving optimal performance. For example, studies using discrimination learning tasks have shown that, both in humans (Lawrence, Sahakian, Rogers, Hodges, & Robbins, 1999; Kalish & Kruschke, 2000) and in rats (Oswald et al., 2001; Sutherland & Mackintosh, 1971), a stimulus feature (e.g. blue colour) that predicts a reinforcer leads to greater attention being allocated to the same feature (blue) or to other features from the same dimension (e.g. colour). The learnt change in attention allocation observed in intra-dimensional shift/extra-dimensional shift experiments generalises between features based on their similarity, resulting in more attention to features from the same dimension than to those from a different dimension (e.g. shape) (Mackintosh, 1975; for review, see Le Pelley, 2004). In contrast, the model by Pearce and Hall (1980) states that cues with uncertain consequences capture the most attention. Because unreliable cues are surprising, they will initially capture attention preferentially, leading to rapid learning about their significance. Several hybrid models have emerged in an attempt to reconcile the influence of predictability and uncertainty (Kakade & Dayan, 2002; for reviews, see Esber & Haselgrove, 2011; Pearce & Mackintosh, 2010). Moreover, typical studies of the influence of attention on memory only present each stimulus once (Christianson, Loftus, Hoffman, & Loftus, 1991; Kim, Vossel, & Gamer, 2013; Riggs et al., 2011; Talmi, Schimmack, Paterson, & Moscovitch, 2007). Our main objective here was to develop a new animal test of the impact of motivational salience on attention, inspired more closely by the human attention literature. For this purpose we will use object preference in a NOR-like task.

There is burgeoning interest in the influence of emotion on human cognition, including the allocation of attention (Golomb, Nguyen-Phuc, Mazer, McCarthy, & Chun, 2010; Pourtois, Schettino, & Vuilleumier, 2013), and how biased attention influences subsequent memory (Ack Baraly, Hot, Davidson, & Talmi, 2017). Yet the term ‘emotion’ is notoriously nebulous, even in humans (Izard, 2010). For comparative purposes, it is now considered more appropriate to focus research on survival-related functions (LeDoux, 2012). Motivational salience is a latent variable that indicates the reinforcing potential of a stimulus, and translates well across species (Schultz, 2016). Indeed, stimuli that are thought to evoke emotion in animals are those that predict fitness-relevant consequences – either reward or punishment (Rolls, 2000). Therefore, in order to develop a rodent test of emotional biases in attention that could be useful across species, we chose to operationalise emotion by manipulating the motivational salience of objects. There is a solid basis for this approach in human research too, with evidence that humans allocate preferential attention to stimuli that predict reward or punishment (Le Pelley et al., 2016). The influence of motivational salience on either attention or memory, as they are measured in the NOR task, has not been investigated, but could be of considerable interest. The structure of the NOR task suggests that in order to do so, we would need to manipulate the motivational salience of objects. Here we manipulate the motivational salience of objects in a NOR-like task with a focus on the impact of this manipulation on attention, quantified through object preference.

Evidence for the influence of motivational significance on human attention in sensory domains outside vision is sparse, despite abundant research on different sensory modalities involved in bottom-up and top-down attention processes (Spence, 2010). While humans are predominantly influenced by visual stimuli (Shapiro, Egerman, & Klein, 1984), most mammals, including rodents, rely mainly on odours and somatosensation to obtain information about their environment (Brennan & Kendrick, 2006; Johnston, 2003). Odours from conspecifics of opposite sex are thought to carry positive reward values, because they elicit an approach response to promote reproductive behaviours (Beny & Kimchi, 2014). According to Huckins et al. (2013), conspecific urine smell is a ‘social odour’ and, therefore, high in motivational significance.

The motivational significance of odours bearing reproductive value does not seem to depend on the animal’s prior sexual experience. This is evident in the laboratory, where sexually inexperienced male rodents display typical sexual behaviours when exposed to female odour (Beny & Kimchi, 2014). Given the socio-biological relevance of female odour and its known motivational effects on animals, we decided to manipulate the motivational salience of objects by labelling some of these with female odour. To do so, we placed the objects for some hours in bedding taken from cages that housed female mice (see methods, below). Henceforth, we refer to these objects as O+ objects. Previous research used social odours to examine olfactory processing (Arbuckle, Smith, Gomez, & Lugo, 2015; Rattazzi et al., 2015; Yang & Crawley, 2009) but provides limited information on how to best utilise odour to change the behaviour of mice in the NOR task. The experiments reported here were designed to clarify the impact of O+ objects on male mice within the specific experimental context of the NOR.

Our study is the first to test the influence of motivational salience on attention in the widely-used NOR task. In this study we focused on testing whether mice pay more attention to objects labelled with female odour, and on quantifying the attentional resources that they allocate to these. In future, this work can help establish a reliable protocol for behavioural and neurobiological investigations of the influence of motivational salience on memory, as it is measured through the NOR and its more sophisticated variants, such as the what-where-which task in mouse and rat (Davis, Eacott, Easton, & Gigg, 2013; Eacott & Norman, 2004).

## Methods

Seven experiments were designed to quantify the extent of preferential attention in male mice to motivationally-salient objects. Attention was operationalised as exploration time of objects placed in a Y-maze. In each experimental trial we compared attention to the experimental O+ object, versus control objects: either copies of the same objects that were labelled with a non-social odour (almond extract, O^a^) or no odour (O-), or to novel objects (with no odour, X-), which were placed in either novel or familiar locations.

### Animals

Experiments 1-6 were performed using eight adult male C57BL/6J mice (Charles River, UK), which were 14 weeks (3.5 months) old at the start of the experiments. These experiments were completed within a period of two months, at a chronological order that, due to experimental constraints, slightly differed from their numbering here (Experiment 2 was carried out after Experiment 4). Note that this strain of mice have a stable performance on the NOR at least until they are 12 months old (Traschütz, Kummer, Schwartz, & Heneka, 2018).

Experiment 7 was conducted after Experiment 6, with a new group of eight adult male C57BL/6J mice (Charles River, UK), which were 14 weeks (3.5 months) old at the start of the experiment.

Mice were maintained by the Biological Services Facility (BSF), University of Manchester, UK and housed in groups of four individuals in ventilated Techniplast cages, in standard conditions (20°± 2°C temperature and 55±5% humidity) on a 12:12 light/dark cycle, with *ad libitum* access to food and water. All experiments took place between 9:30am-12:30pm; while there are no known circadian influences on NOR in this strain, and we did not see differences between the performance of animals from this strain in an operant conditioning task, we cannot rule out that the results may be different if animals were tested at another time of day. All mice were ear-punched for identification. All procedures were conducted in conformity with the University of Manchester BSF regulations for animal husbandry and the Home Office Animals (Scientific Procedures) Act (1986) and were approved by University of Manchester Ethical Review Panel under a project licenced granted by the UK Home Office.

### Apparatus

Experiments were performed in a row of four custom Y-mazes with three identical white, opaque plastic arms, (length 160mm, height 280mm, internal width 4800mm) diverging at a 120° angle from each other (designed by Jack Rivers-Auty and constructed by Plastic Formers Ltd, UK). Each arm became wider at the end to form a small square arena (length 92mm x width 90mm). The square arenas could be differentiated by the presence of salient visual cues on the walls. Individual mice were randomly assigned to a particular Y-maze throughout habituation and testing. Mice were always released inside the middle arm (the arm closest to the nearest room wall), facing away from arms that contained objects (see Materials). This strategy, following the guidelines by Antunes and Biala (2012) and ensured that external pressure to explore the objects was avoided. During the interval between exposures in the habituation stage and the inter-trial intervals of the experimental task, mice from each cage group were placed in separate holding cages (standard housing cages, Techniplast), which remained the same throughout the project. Mice from each cage group were run simultaneously in the four Y-mazes placed next to each other. Video cameras (JVC, 40x optical zoom) placed above the Y-mazes recorded animal performance.

## Materials

### Objects

These were built either from LEGO^®^ pieces or various other plastic shapes such as plastic eggs halves (Figure 1). Because the eggs were identical in shape, the difference in their colour as well as their position in the maze (face-down or on one side with either convex or concave side facing the animal) were used to create different-looking objects. All objects were odour-neutral and made of plastic in order to avoid material preference, minimise odour saturation and facilitate cleaning. The objects were attached to the floor inside the Y-maze with Blu-Tack®. Objects used in the habituation phase were plastic letters and were never used in subsequent experimental trials. All objects were unique, in order to avoid habituation.

**Figure 1:**
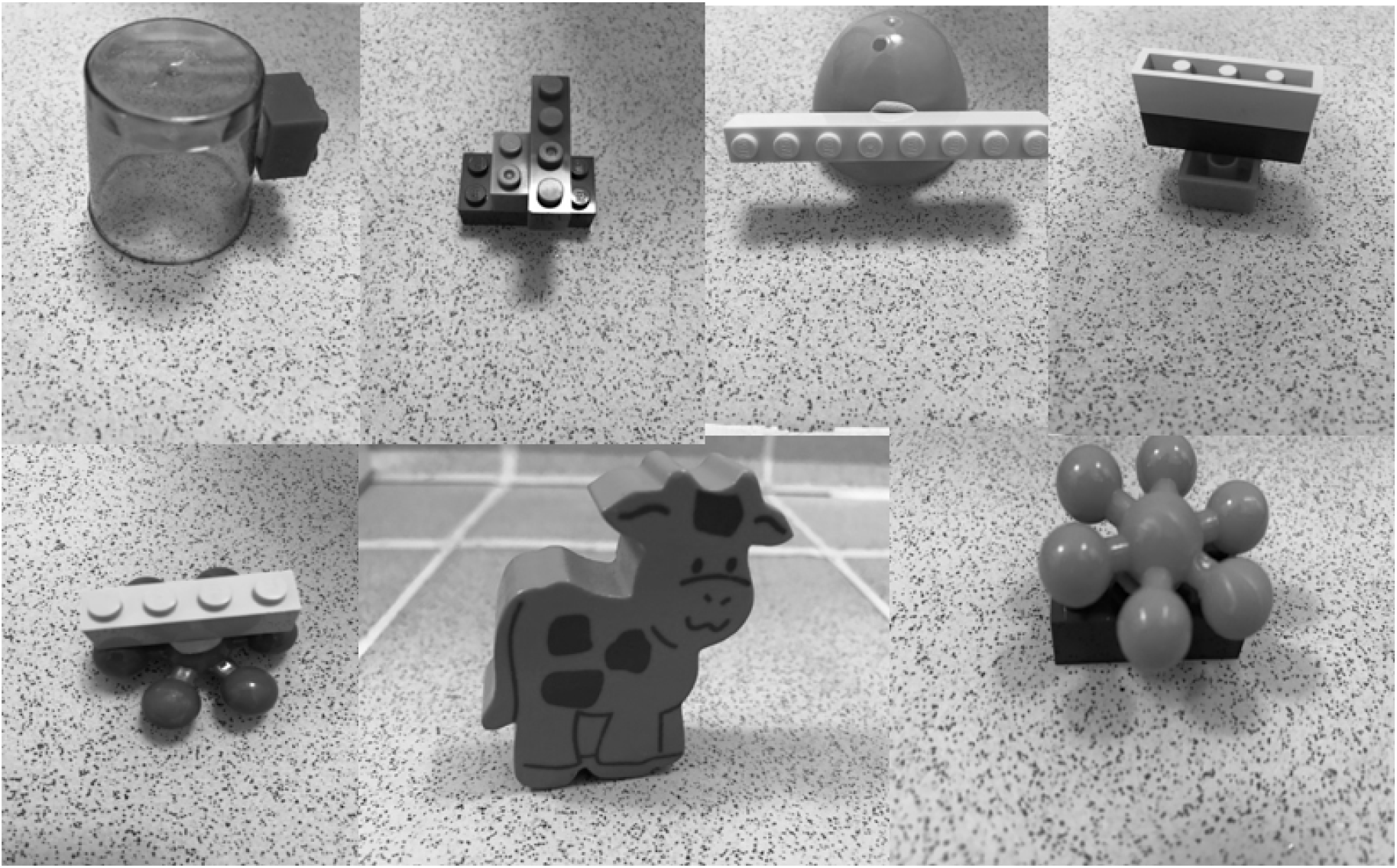
Examples for objects used in the study.

Each animal was exposed to a given object only in one experimental trial across the entire study (once across experiments 1-7). This means that the number of objects an animal experienced overall in each experiment equates to the number of trials in each experiment, listed in the procedure, below. However, in experiments 2-7, where in each experimental trial animals were exposed to the maze twice, several copies of the same object were used. The particular object used in a given trial, the order in which objects were presented and their locations in the maze (left or right choice arm) were counterbalanced across animals in each experiment. Apart from Experiment 1, each animal was exposed only to one O+ on each day; whether this was in the first or second trial of the day was counterbalanced.

### Female odour

In this study we opted for a pragmatic approach to obtain a motivationally-salient odour. For each experiment, soiled cage bedding from cages containing four female mice (C57BL/6J strain) was used to label objects with female odour. The objects to be so “labelled” were placed in the bedding the day prior to each trial, around midday (12:00pm). For the experiments that lasted four days (Experiments 1, 3 and 4), in order to prevent odour habituation, the bedding with female odour was replaced with bedding from a cage containing a different group of females of the same strain halfway through each experiment (at the end of day 2). For experiments that lasted only 2 days (Experiments 2, 5, 6 and 7), the bedding remained the same throughout each experiment, but was changed before the start of the next experiment. Y-mazes and objects were cleaned thoroughly with 70% ethanol and wiped with paper towels between trials with mice from different group cages, as well as at the end of each daily session.

### Control odour

Compound odour mixtures, such as a flower, nut or fruit, are prevalent in behavioural studies of mice (Arbuckle, Smith, Gomez, & Lugo, 2015; Rattazzi, Cariboni, Poojara, Shenfeld, & D’Acquisto, 2015; Yang & Crawley, 2010). Because female odour is also a compound odour mixture, we decided to use another compound mixture here as a control odour. We selected almond odour, one of the odours used by Rattazzi et al. (2015). The odour of almond extract is mildly motivationally significant, since it is from a natural food source but distinct from the food laboratory mice are used to (Huckins, Logan, & Sanchez-Andrade, 2013). Hutckins and colleagues used almond odour as a non-social odour alongside social odours in rodent experiments investigating odour-mediated behaviour and odour identification ability.

To label the filter paper with almond odour we dipped a cotton-tipped applicator in pure almond oil (100% concentration, by Atlantic Aromatics, Bray, Co. Wicklow, Republic of Ireland) and then scrubbed it on a piece of filter paper (4cm diameter). To label O^a^ objects with almond odour, we scrubbed O objects with cotton-tipped applicators dipped in almond oil. Then, in order to remove the oiliness from these objects which could have led to a possible bias in attention capture (arising from a difference in texture between O+ and O-), O^a^ objects were also gently wiped with a paper towel before being placed in the Y-mazes.

## Procedure

### Overview

The tasks used in this study were fashioned after the classical NOR task (Berlyne, 1950). While it is well known that rodents explore preferentially objects that are novel, or placed in novel locations (Aggleton, Albasser, Aggleton, Poirier, & Pearce, 2010; Barker & Warburton, 2011; Ennaceur, Neave, & Aggleton, 1997), here we marry this task with the literature on preferential approach to female odour (Connor, 1972; Mackintosh, 1970) in order to quantify the relative potency of female odour, versus other salient object features, to bias exploration of objects in mice.

The first two experiments assessed the ability of a single object to capture attention in mice. This object was either O+ or O- and either novel (Experiment 1) or familiar (Experiment 2). We hypothesised that because the motivational salience of O+ was greater than that of O-, the former would attract more attention in both experiments.

Next, we asked how much attention would O+ attract, if it were not only motivationally salient, but also placed at either a novel (Experiment 3) or a familiar (Experiment 4) location. Novel location is known to capture attention (Ennaceur, Neave, & Aggleton, 1997) and, thus, it was particularly interesting to study whether female odour would capture even more attention. We hypothesised that in Experiment 3, when the location of O+ was novel, O+ would capture more attention than O-, because the former benefited from the advantage of two salient features, namely, motivational salience and novel location. However, in Experiment 4 it was harder to predict whether O+ would still ‘win’ over O-, due to the fact that the novel location of the latter was now competing with the odour of the former for attention.

In the next two experiments, we used another well-known object feature, novel object identity, and assessed how strongly attention would be captured by female odour compared to a novel object (X-). Novel objects are known to recruit more attention than familiar objects in the traditional NOR task. Therefore, we wanted to know how attention to O+ would be influenced by the presence of X-. Experiment 5 tested the attention captured by O+ at a novel location versus X- at a familiar location. Here, O+ had the advantage of motivational salience and location novelty, while X- had the advantage of novel identity. In Experiment 6, X- was placed at a novel location and O+ at a familiar location. As X- here had the advantage of both novel identity and novel location, the question was whether it would attract more attention than O+.

In the final experiment we addressed the fact that in Experiments 1-6 we compared an odour stimulus with visual novelty. Experiment 7 set out to compare female odour to another odour to clarify whether female odour is particularly motivationally salient within the context of our modified NOR. Such finding will be consistent with preliminary evidence that social odour increases sniffing time in male mice compared to other social and non-social odours (Rattazzi, Cariboni, Poojara, Shoenfeld, & D’Acquisto, 2015). We predicted that despite the mild motivational significance of almond odour, animals in Experiment 7 will allocate more attention to female odour.

### Habituation

Before the study began animals were handled for one minute on two consecutive days so that they became accustomed to the experimenter. Next, animals were habituated to the testing apparatus and objects over a five-day period. The first day of habituation consisted of a 10-minute cage group habituation session to the Y-mazes, in which mice from a given cage were placed together in one Y-maze, in the absence of objects. On the following day, the mice were exposed individually to the Y-maze for 10 minutes, again without objects. On the third day, the same steps from day 2 were repeated, but this time an O was present in either the left or right choice arm. On the fourth day, following the same steps as on day 3, each animal was exposed to another O-, but in the opposite arm to the one on day 3. The last day of habituation consisted of two exposures, this time with an O+ in either the left or right arm of the maze. The duration of exposure was 10 minutes and the interval between exposures was 5 minutes. During the 5-minute interval, the mice were placed in their holding cages and the O+ presented in the first exposure was replaced with different O+ for the second exposure. Prior to Experiment 7, mice were habituated to O+ following the same procedure as in Experiment 1-6, and in addition they were habituated to O^a^ objects in either left or right arm.

### Experimental task

Experiments 1, 3 and 4 were conducted on four consecutive days. Experiments 2, 5, 6, 7 (and pre-test for 7) were conducted on two consecutive days. Figure 2 depicts what experimental trials looked like in each experiment. In Experiment 1 the experimental trial included only a single exposure to the maze. In subsequent experiments, each experimental trial included two exposures to the maze. The second exposure came immediately after the first, with no time delay. In all experiments, each day of testing included two experimental trials. The interval between experimental trials was 5 minutes. During that time, animals were placed in holding cages and the objects replaced for the next trial.

**Figure 2:**
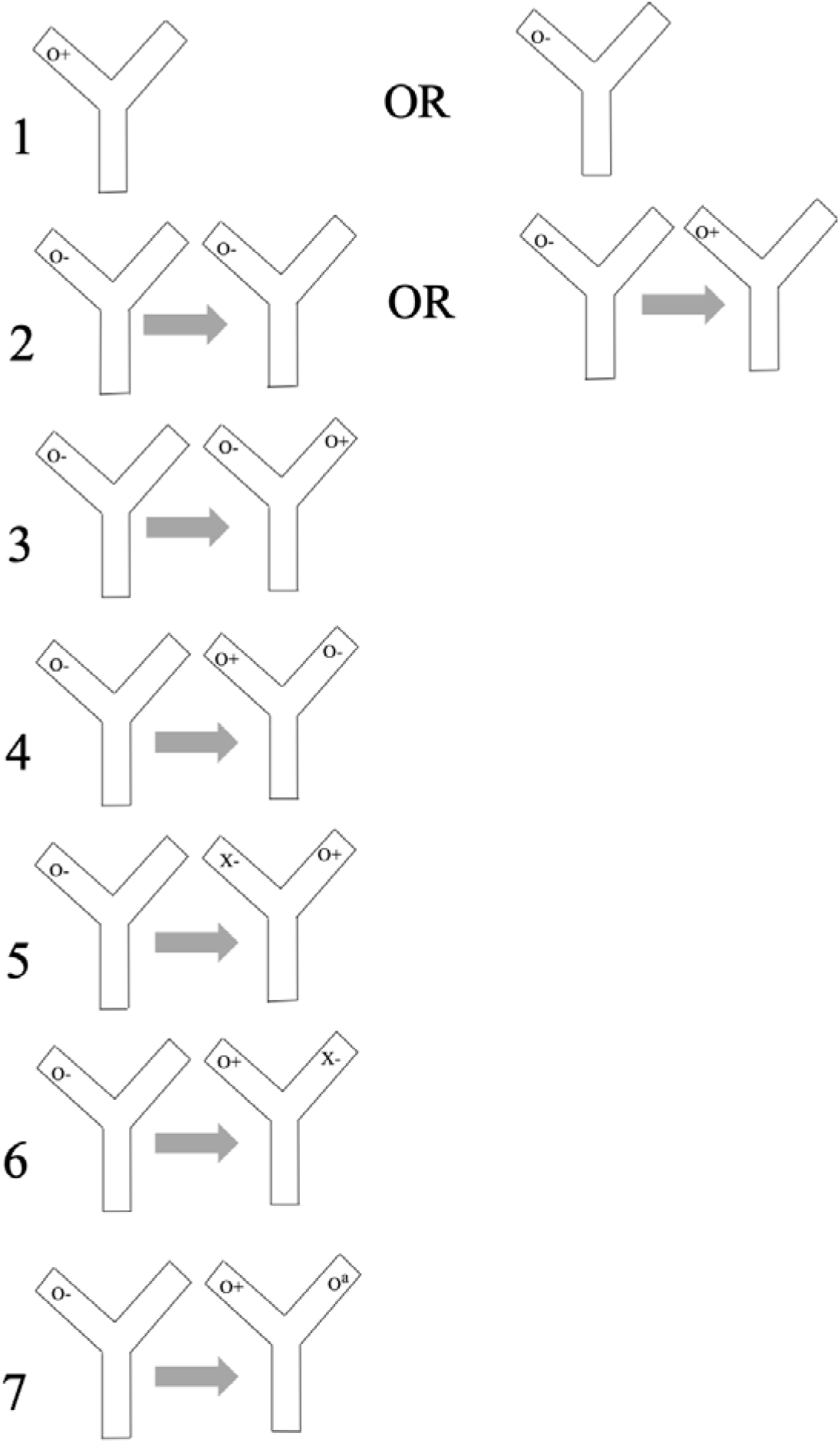
A graphical depiction of the structure of experimental trials in Experiments 1-7. Experiments are presented separately in each row, denoted with a number. All conditions were fully counterbalanced, including the location of the O- object in the first exposure of the animal to the arena, but in the figure that object is always placed in the left arm. O+: Objects labelled with female odour O-: Objects that are identical in identity to O+, but which were not labelled with any odour. O^a^: Objects that are identical in identity to O+, but which were labelled with almond odour. X-: Objects which were different in identity to O+, O-, or O^a^. They were not labelled with any odour.

#### Experiment 1

This experiment consisted of eight trials in total per animal. Trials included one exposure to a single object, either O+ or O-. The duration of each trial was one minute (see row 1 of Fig. 2).

#### Experiment 2

This experiment was composed of four trials per animal. Each trial included an initial exposure to O- for one minute (‘familiarisation’), either in the left or the right arm of the maze, followed by an exposure either to O- or O+ for 3 minutes (‘test’; see row 2 of Fig. 2).

#### Experiment 3

This experiment included eight trials in total per animal. Each trial included a familiarisation phase, where animals were exposed to O- for one minute, either in the left or the right arm of the maze. The location of O- in the familiarisation phase is referred to as the ‘familiar’ location. The other location – the arm that was empty during the familiarisation phase – is referred to as the ‘novel’ location. During the test phase animals were exposed to both O- and O+ for 3 minutes. The O+ was always placed in the ‘novel’ location (see row 3 of Fig. 2).

#### Experiment 4

This experiment was identical to experiment 2, except that in the test phase O+ was always placed in a familiar location (see row 4 of Fig. 2).

#### Experiment 5

This experiment was composed of four trials in total per animal. Each trial included a familiarisation phase, where animals were exposed to O- for one minute, either in the left or the right arm of the maze. The location of O- in the familiarisation phase is referred to as the ‘familiar’ location. The other location – the arm that was empty during the familiarisation phase – is referred to as the ‘novel’ location. During the test phase animals were exposed to both O+ and X- (an entirely novel object) for 3 minutes. The O+ was always placed in the ‘novel’ location (see row 5 of Fig. 2).

#### Experiment 6

This experiment was identical to experiment 5, except that O+ in the test phase always occupied a familiar location (see row 6 of Fig. 2).

#### Pre-test for experiment 7

In order to ensure that the mice were to some extent attracted to almond odour, we tested the preference of mice for almond odour. This experiment was composed of four trials in total per animal. Animals were simultaneously exposed to two filter papers, one without any odour and the other labelled with almond odour, at randomly determined locations in the Y-maze (left or right arm).

#### Experiment 7

In this experiment, O+ in the test phase was always placed at a familiar location and the O^a^ occupied a novel location (see row 7 of Fig. 2). The number of trials and the way the experiment was conducted was identical to Experiment 6.

### Data analysis

Exploratory behaviour was recorded with video cameras. Subsequently, object exploration time for each object was scored using the Novel Object Timer software (Jack Rivers-Auty; Novel Object Recognition Task Timer, 2015). An animal was considered to be exploring an object when its nose was within 2cm of the object and directed at the object. Sitting next to the object or climbing on top of the object was not regarded as exploratory behaviour. Each trial was scored twice for accuracy and the average of the two scorings taken as the object exploration time per trial. The mean exploration time of O+, O-, O^a^ and X- objects was obtained by averaging all trials with these objects for each animal.

For Experiments 3-7, where during their second exposure to the arena animals were presented with two objects, the discrimination index (D2, Antunes & Biala, 2012) was used to assess preferential attention. D2) was calculated using the formula: D2 = (T_O+_ – T_Other_) / (T_O+_ + T_Other_), where *T_O+_* is the mean exploration time of O+ objects and *T_Other_* is the mean exploration time of either O-, O^a^ or X- objects. The values for this index are bound within a range of −1 to 1; positive D2 values indicated preference for O+, while negative values signified preference for other objects, and a value of 0 signified no preference (Burke, Wallace, Nematollahi, Uprety, & Barnes, 2010; Oliveira, Hawk, Abel, & Havekes, 2010).

Experiments 1, 3 and 4 (which were conducted consecutively, in terms of their chronological order) were completed over 4 days. Initially, we did not have a specific hypothesis for an effect of experimental day, and consequently, the counterbalancing of Experiment 1 did not take the ‘day’ into account. Therefore, we averaged across days, and analysed D2 values in that experiment with a two-tailed paired t-test, comparing O+ to O-. When the raw data indicated a potential habituation of the preference for O+ across days, we took ‘day’ into account in the counterbalancing of Experiments 3-4, to be able to investigate the effect of ‘day’ formally. Therefore, these experiments were analysed with a repeated-measures ANOVA with two within-subject factors: Object Type (O+, O-) and Experiment Day (day 1-4). As we did not find a significant main effect or interactions with Experiment Day, we did not examine it in the following experiments. Thus, in Experiments 2 and 5-7 the differences between Object Types were again analysed with two-tailed paired t-tests. These tests always compared the exploration of the experimental object, O+, to the exploration of the control object (see Figure 2), which was either O- (Experiment 2), X- (Experiments 5-6) or O^a^ (Experiment 7). Statistical tests were performed in GraphPad Prism 8.

## Results

### Experiment 1: Novel objects O+ vs. O-

Two animals were excluded from the analysis of Experiment 1 due to a counterbalancing error; including them in the sample did not change the results significantly. The total time animals spent exploring O+ was significantly higher than the total exploration time for O-, *t*(5)=4.11, *p*=0.0092, as illustrated in Figure 3 (left panel). This finding demonstrates that when both O+ and O- are novel, the former attracts more attention.

**Figure 3:**
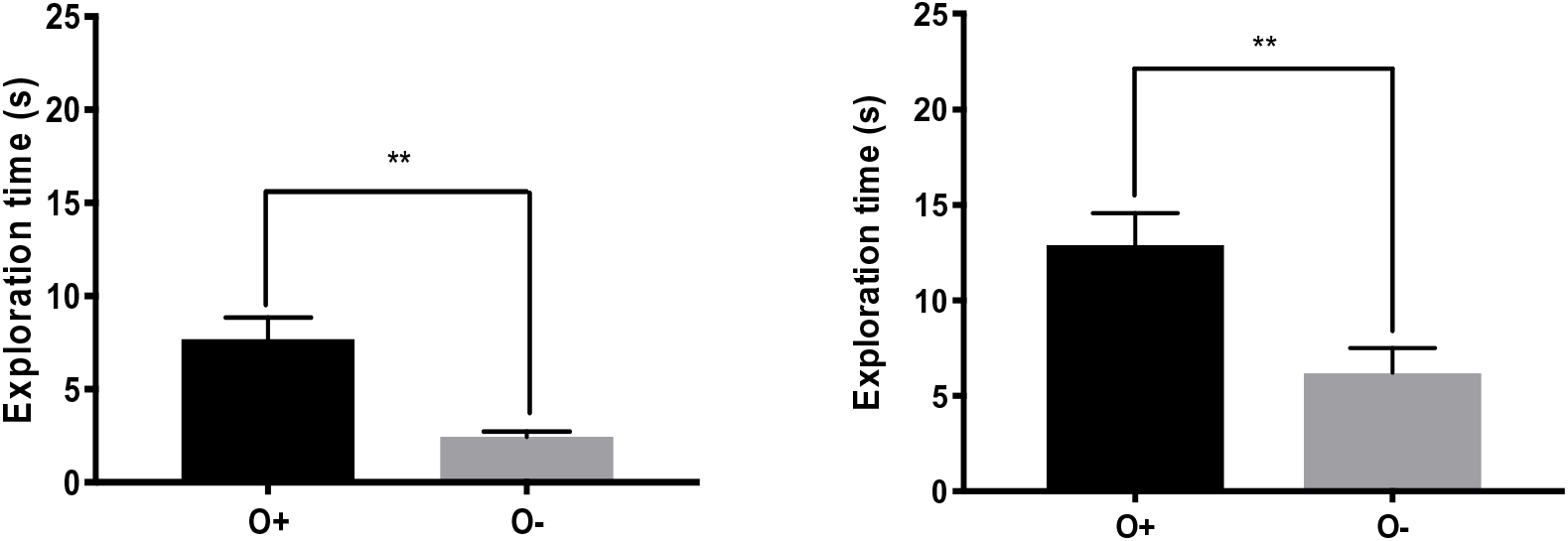
Exploration of single objects in Experiments 1 and 2. Bar graphs represent object exploration times (in seconds) averaged across all animals (6 in Experiment 1 and 8 in Experiment 2). Data are presented as means ± SEM. Statistical significance is expressed with ** (p≤0.01). **Left:** Results of Experiment 1: Novel O+ vs. Novel O-. **Right:** Results of Experiment 2: Familiar O+ vs. Familiar O-.

### Experiment 2: Familiar objects O+ vs. O-

The total object exploration time in Experiment 2 (Fig. 3 right panel) was significantly higher for O+ than O-, t(7)=3.67, p=0.008. This finding demonstrates that when both O+ and O- are familiar, the former attracts more attention.

### Experiment 3: Competition between O+ in a novel location and O- in a familiar location

Figure 4 shows that the total exploration time for O+ was significantly higher than that for O-. Object exploration time was analysed across experimental days for any effect of day or interaction between object type and day. A repeated-measures two-way-ANOVA test found that there was a significant effect of object type, F(1,7)=47.84, p<0.001. While numerically this difference diminished across days, neither the effect of day, F(3,21)=2.93, p=0.06, nor the interaction, F(3,21)<1, p= 0.53, were significant. Animals demonstrated a significant preference for O+ over O-, as indicated by the mean D2 value, which was significantly higher than zero, t(7)=9.33, p<0.0001.

**Figure 4:**
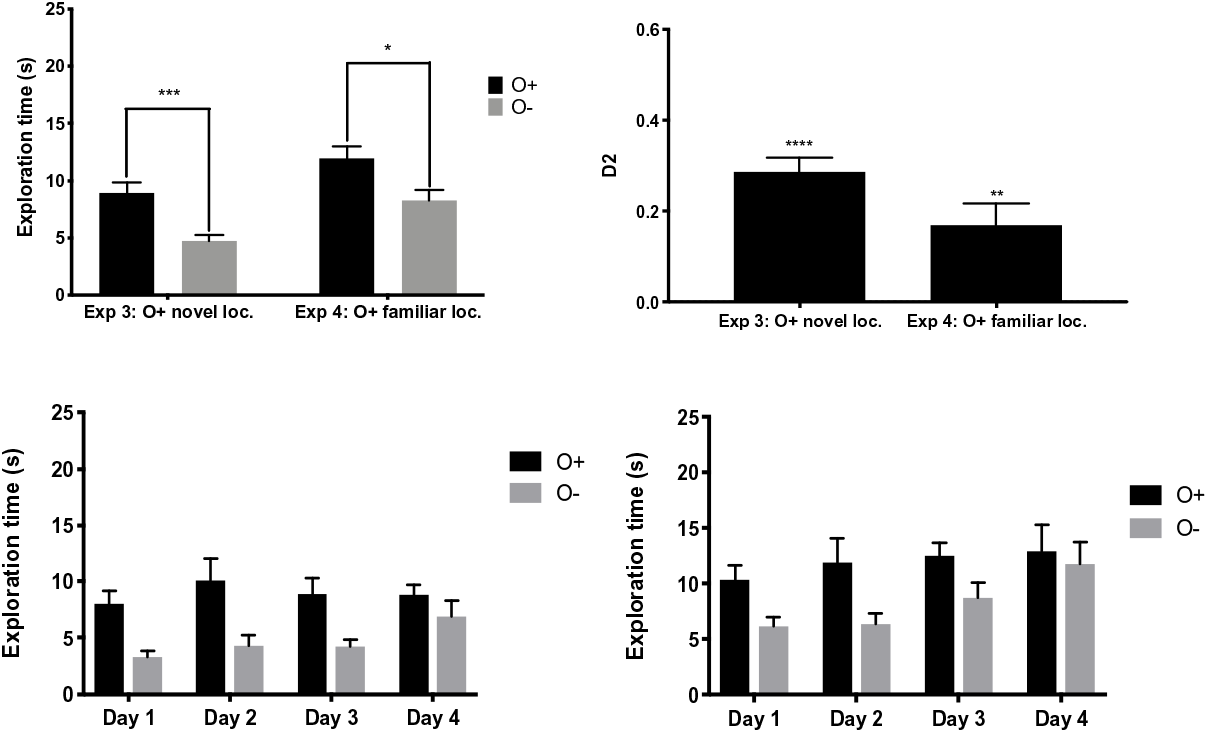
Results of Experiments 3 and 4. Means of object exploration time (seconds) were calculated by averaging across all 8 animals. Bar graphs represent average exploration times (in seconds). Mean D2 values represent the average of the D2 values. Data are presented as means ± SEM. Statistical significance is expressed as * (p≤0.05), *** (p≤0.001) or **** (p≤0.0001). **Top left:** Object exploration time in Experiments 3 and 4. **Top right:** Effect of location familiarity on odour preference. The mean D2 value was significantly greater than zero in both experiments. **Bottom:** Daily object exploration in Experiment 3 (Left) and 4 (Right).

### Experiment 4: Competition between O+ in a familiar location and O- in a novel location

The total exploration time for O+ was higher than that for O- (Figure 4), and the average D2 value again indicated a preference for O+ over O-, because it was significantly different to zero, t(7)=3.67, p=0.008, and positive. Object exploration time was analysed across experimental days for any day effect or interaction between object type and day. Replicating the above results, there was a significant effect of object type, F(1,7)=12.02, p=0.01. As in the previous experiment, here the effect of day (F(3,21)=2.24, p=0.11) and the interaction (F(3,21)=1.13, p=0.36) were not significant.

### Experiment 5: Competition between O+ in a novel location and X- in a familiar location

As shown in Figure 5, the total object exploration time in Experiment 5 was significantly higher for O+ than for X (t(7)=5.63, p=0.0008). A one sample t-test showed that the D2 value was significantly different to zero, indicating a preference for O+ over X (t(7)=4.92, p=0.0017).

**Figure 5:**
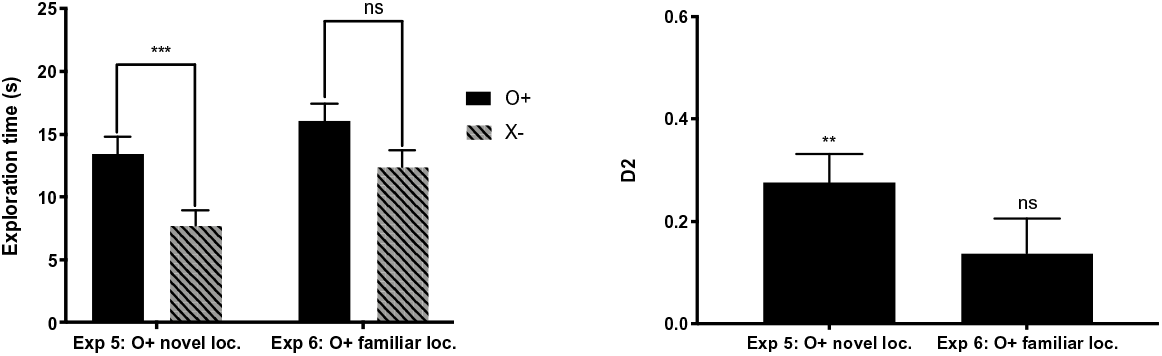
Results of Experiments 5 and 6. Data are presented as means ± SEM across 8 animals. Statistical significance is expressed as ns (p>0.05), ** (p≤0.01) or *** (p≤0.001). **Left:** Effect of object identity and location familiarity on exploration time. Bar graphs represent average object exploration times (in seconds). **Right:** Effect of object identity and location familiarity on object preference. Mean D2, the average of the D2 values, was significantly greater than zero in Experiment 5 but not in Experiment 6.

### Experiment 6: Competition between O+ in a familiar location and X- in a novel location

The total object exploration time in Experiment 6 was higher for O+ than for X-, however, the difference was not statistically significant (t(7)=1.8, p=0.1158, Figure 5). The D2 value in this experiment was also not significantly different to zero, suggesting no significant preference for O+ over X- (t(7)=2.023, p=0.0828).

### Analysis across Experiments 5 and 6

Since there were no differences in the duration of exploration of O+ and X- objects in Experiment 6, we determined whether the data holds other clues to animal preference. One possibility was that animals indicated a subtle preference by exploring a particular object type first; this is referred to as *first object choice*. A binomial test revealed that there was not enough evidence to reject the null hypothesis, according to which animals were equally likely to explore either object, (p=0.1402 in Experiment 5, p=0.2153 in Experiment 6).

### Pre-test for experiment 7: Almond oil as a salient stimulus

It was necessary to investigate first whether mice are able to detect the almond odour and, moreover, if they found this smell attractive. Figure 6 shows that the total exploration time of filter papers with almond odour was significantly greater than the exploration time of an odour-free filter paper, t(7)=2.9, p=0.0229. The D2 value, which was found to be significantly different to zero, t(7)=3.5, p=0.0101, confirms the preference of mice for almond odour compared to filter paper alone.

**Figure 6:**
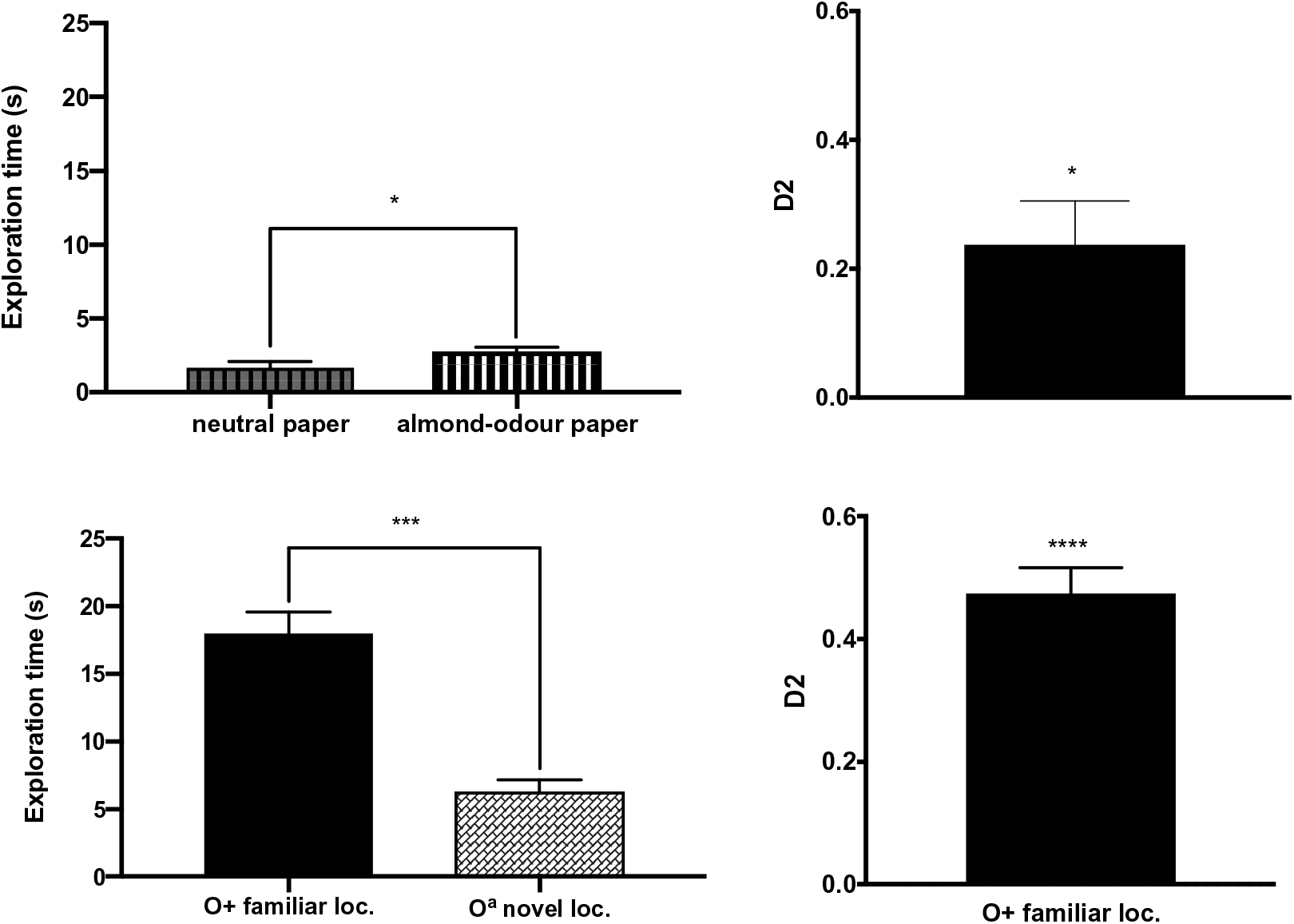
Results of Experiment 7. Means of object exploration time (seconds) were calculated by averaging across all 8 animals. Bar graphs represent average exploration times (in seconds). Mean D2 values represent the average of the D2 values. Data are presented as means ± SEM. Statistical significance is expressed as * (p≤0.05), *** (p≤0.001) or **** (p≤0.0001). **Top left:** Effect of odour on exploration time in pre-test for experiment 7. **Top right:** Odour preference in pre-test for experiment 7. The mean D2 value was significantly greater than zero. **Bottom left:** Effect of odour on exploration time in Experiment 7. **Bottom right:** Effect of odour on object preference in Experiment 7. The mean D2 value was significantly greater than zero.

### Experiment 7: Competition between O+ in a familiar location and O^a^ in a novel location

The overall exploration time for O+ was significantly higher than that for O^a^, t(7)=7.5, p=0.0001, and the mean D2 value was significantly different to zero, t(7)=11.5, p<0.0001 (Figure 6), indicating preference for O+ over O^a^.

One explanation that could have accounted for the very high preference of mice for O+ compared with O^a^ was that, in fact, the high concentration of almond oil resulted in the odour of O^a^ being aversive to the mice. According to Saraiva et al. (2016), some odours that are attractive at a low concentration can become aversive in high concentrations. In Experiment 7, however, we were able to determine whether the mice were attracted or repulsed by the almond smell by looking at the video recordings. Mice did not display avoidance behaviour toward O^a^ objects; they not only sniffed the O^a^ objects multiple times upon their first discovery of such an object in one of the Y-maze’s arms, but they also returned to this object several times during the 3-minute test. Finally, the results of the pre-test allows us to conclude that the mice preferred the almond odour to the odour-free paper, clarifying that it was not aversive.

## Discussion

The amount of attention that C57BL/6J male mice allocate to objects was measured here through the duration over which they explored single objects (Experiments 1-2), and the preference they exhibited to explore one object over another (Experiments 3-7). We found that these mice attended objects labelled with a female odour more than objects that were not labelled with an odour, and also more than objects labelled with a non-social odour. Additionally, they attended objects with female odour more than non-odour novel objects placed in familiar locations, and more than non-odour familiar objects that were placed in novel locations. They attended objects with female odour approximately as much as non-odour novel objects that were placed in novel locations.

It is known that female odour is a primary reinforcer for male mice. The only experience that these laboratory mice had with female odours was their mother’s smell during weaning, after which they were isolated from female conspecifics. Previous studies have shown that female odour represents a positive arousing stimulus for laboratory male mice, and that sexually-inexperienced (naïve) male mice display sexual behaviours towards female conspecifics and manifest aggression towards other male mice (Beny & Kimchi, 2014; Connor, 1972; Mackintosh, 1970). Yet, despite these well-known effects of female odour, we could not be sure, before we conducted the present experiments, whether we would be able to demonstrate a measureable bias towards objects that were ‘labelled’ O+ by storing them for some hours in bedding taken from cages that housed female mice. Previous research has shown that it is difficult to know in advance which aspect of a female will elicit approach in males and how approach depends on the sexual experience of males; for example, adult male rats without copulation experience did not approach a taxidermy model of a female (Holloway & Domjan, 1993). Our experimental choices – from the method we used to ‘label’ the objects, the use of the D2 index, and the within-subject nature of each experiment, where we subjected the same animals to up to 8 trials across as many as 4 days – could have influenced the particular behaviour exhibited. The finding that male mice exhibited substantial, robust preference for O+ objects advances a protocol for incorporating a motivationally-salient odour within the NOR task. In future, such protocols can be used to study the influence of motivational salience on memory performance in the NOR and related paradigms. Investigations of working memory have shown that incorporating odour can be translationally productive; for example, mouse models of Alzheimer’s disease show poor odour memory span, as do AD patients (Bahuleyan & Singh, 2012). O+ objects can also be used to study episodic memory in the WWWhich task (Davis et al., 2013; Eacott & Norman, 2004), a state-of-the art task to investigate episodic-like memory in rodents.

In each experimental trial in Experiment 1 animals were exposed to a single object once; it either O+ or O-. Because both objects were novel, the preference of animals to explore O+ in that experiment can be attributed unambiguously to motivational biases. Research on emotional cognition is often satisfied with the demonstration of difference between two categories of stimuli, a nominally emotional one and a nominally neutral one. Here, we endeavoured to go beyond the nominal scale to the ordinal scale and elucidate *how much attention* is allocated to female odour compared to other forms of salience. The experiments were designed to answer this question by creating situations where this odour can compete for attention with objects expressing other salient features.

In order to quantify the degree of preference for the O+, in subsequent experiments we compared their attention to O+ to their attention to a control object. In Experiments 2-7 animals were first introduced to the control stimulus, O-, and then they were presented with both experimental and control stimuli. During the test stage of Experiment 2 animals were presented with a single object: either an O- or an O+. Because they were familiarised with an O- in the same location beforehand, the animals likely recognised the identity, as well as the location, of the object. We can therefore attribute their preference towards O+ to its odour. However, because the lack of odour of the O- was familiar, and the odour of the O+ was novel, Experiment 2 cannot disambiguate whether the attentional bias towards O+ has to do with novelty or with motivational significance of the odour.

In Experiment 7 we compared the degree of attention allocation produced by female odour to that of another olfactory stimulus with a milder motivational significance. We labelled the Oa objects with undiluted almond oil, while Rattazzi, Cariboni, Poojara, Shoenfeld, & D’Acquisto (2015) used a 100-fold dilution of almond extract (10μl almond extract to 990μl distilled water). The pre-test confirmed that this odour was mildly attractive to the mice. Given that the potency of almond oil in our experiment was likely higher than that used by Rattazzi and colleagues, almond odour could have outcompeted female odour in animals’ attention. This was, however, not the case. In the first exposure the mice explored O-, and in the second exposure, they were presented with O+ in half the trials and Oa in the other half. Mice consistently showed significantly higher exploration time and preference for O+ in Experiment 7. This observation not only confirmed that mice are more attracted to female odour than to a novel non-social odour (which was already well-established in the literature), but also that this holds true even when almond odour was intense, and found on an object placed at a novel location in a NOR-like set-up. Because in Experiment 7 both O+ and Oa had novel odours, these results suggest also that the attentional bias to explore O+ stems partly from its motivational salience, not only its novelty. Pragmatically, Experiment 7 suggests that almond extract was mildly motivationally salient, suggesting that it can be employed, in future, as a control odour.

Previous studies have already established that, under normal conditions, adult rats show preference for an object at a novel location compared to an identical object at a previously experienced location (Aggleton, Albasser, Aggleton, Poirier, & Pearce, 2010; Barker & Warburton, 2011; Ennaceur et al., 1997). Therefore, it was not surprising that in Experiment 3, O^+^ at a novel location attracted more attention than an O^-^ at a familiar location, since both its odour and location rendered O^+^ more salient than O^-^. Crucially, in Experiment 4, where O^+^ at a familiar location competed with O^-^ at a novel location, O^+^ still attracted more attention. This finding shows that mice will, at least under some conditions, allocate *more* attention to female odour than to object-in-place (location) novelty.

Once we determined that female odour attracts more attention than location novelty, the next logical step was to compare it to another salient object feature – novel object identity. The perceptual salience of novel identity has been the premise of many studies using the NOR task, in which rodents are expected to pay more attention to the novel rather than the familiar object. Therefore, we exposed animals to one object and then, in the next phase, labelled it with female odour, and allowed novel object identity to compete with female odour for attention. In Experiment 5 the salience of O^+^ was due both to its odour and its novel location, while the salience of X^-^ was due to its novel identity. By contrast, in Experiment 6 O+ was salient only because of its odour, while X- was salient because of its novel identity as well as its novel location. We found that more attention was allocated to female odour only when O^+^ was placed in a novel location, not when placed in a familiar location.

Taken together, the results of Experiments 3-6 yield a rough quantification of the potency of female odour for C57BL/6J male mice: objects with this odour attract more attention than novel objects or familiar objects in novel locations, and they attract approximately the same amount of attention as novel objects in novel locations. At present, these conclusions only pertain to the specific ways that we have operationalised motivational salience, object identity, and object location. Further research will be needed to decide whether motivational salience is *generally* equal in magnitude to the salience of novelty in identity and location, and whether these object properties, which here stemmed from multiple modalities (olfaction and vision), combine in an additive or sub-additive manner (Anderson, 2016). The results of Experiments 3-6 confirm that despite the adaptive significance of female odour to male mice, its motivational salience did not completely outcompete that of visual object features. Therefore, in principle, there is a combination of visual object features that could render them equally salient. While here we cannot compare across experiments, it would be interesting to see whether odour can act not only on its own, but also as a component of a stimulating context to modify the degree of attention allocation.

We used the same animals in Experiments 1-6, a choice which risks potential carry-over effects across experiments. In future research it would be beneficial to use new animals in each experiment. Yet it is difficult to see how this factor alone could have produced the set of present results. Notably, animals from the strain perform well on the NOR until they are at least 12 months old (Traschütz et al., 2018), so age alone cannot explain our results. Although it is possible in principle that the odour would become less attractive across experiments, numerically animals explored the O+ more in Experiment 4 than in Experiment 3, and more in Experiment 6 than in Experiment 5, arguing against a habituation account for the results. Still, the null effect in Experiment 6 should be replicated to ensure that it is not a result of habituation to the odour. Using the same animals across experiments 1-6 rendered comparisons across experiments quite tricky, due to potential order effects., so we have refrained from analysing these differences statistically. There was also a benefit in using the same animals across experiments: in Experiments 2-6, animals were somewhat habituated to the female odour, so its novelty perhaps played less of a role. While Rattazzi, Cariboni, Poojara, Shoenfeld, & D’Acquisto (2015) reported that mice habituated to odours after the first exposure, here mice maintained an increasing preference for O+ from Day 1 to Day 4 in Experiments 3 and 4, despite a numerical trend commensurate with slight habituation. It is possible that our results differed from that of Rattazzi and colleagues because we used odour from different females in days 1-2 and in days 3-4. These aspects of our paradigm and results support the claim that attentional bias to O+ was due to its motivational salience, not its novelty.

Our experiments demonstrate conclusively that it is feasible to bias attention in C57BL/6J male mice to objects that were stored with the beddings from female cages, but they are uninformative as to the critical compound within this complex odour. Connor (1972) demonstrated that pheromones found in urine are responsible for dimorphic behaviours. In his experiments, male mice displayed milder aggressive behaviours towards male intruders smelling of female urine and behaved aggressively towards females swabbed with male urine. These behaviours are genetically determined and, therefore, do not require prior learning. However, given the practical method we used to label O+ objects here, we cannot be sure that the biases we observed in attention were due to the odour of female urine, or indeed exactly which compound was responsible and how it influenced endocrine and brain function. Indeed, because we have not controlled the reproductive cycles of females in the cages from which the beddings were taken, it is possible that the attractiveness of the odour to male mice differed across experiments, although note that a mixture of cycles is likely, since virgin females do not synchronise their cycle. Future research could examine whether similar results could be obtained with other odours, such as the odours of male conspecifics. In their study on the effects of odour preferences on rats’ discrimination learning, Devore, Lee, & Linster (2013) classified 53 monomolecular odorants as *high*, *neutral* and *low*, in terms of spontaneous odour preferences. Future work may follow them in establishing an order of preference for mixture of compounds that are common in research with rodents. Additionally, while we can be certain that the attentional effect we observed was not due to a learned association with reproduction, given that the animals were sexually inexperienced, it is possible that the effectiveness of this odour was due to the limited experience that mice had with their mothers as pups. In the future, it would also be useful to compare the degree of attention allocation to female odours with that of odours without primary reinforcing properties that are predictive of a rewarding outcome. Despite these caveats, our work has successfully established female odour as a potent modulator of attention in a modified NOR task, paving the way for future uses in investigations of emotional cognition.

In summary, the present study used object exploration measures to quantify the degree of attention allocation by C57BL/6J male mice to object novelty and female odour. Initial experiments demonstrated that objects with female odours attract more attention than odour-neutral objects. The study further demonstrated that odour can be more potent than location novelty or object identity in the degree of attention allocation, and hinted that the modulatory effect of the odour may combine with that of location or object novelty. Finally, female odour attracted more attention than a mildly attractive non-social odour. These experiments contribute to the understanding of the effects of female odour on attention and provide a reliable protocol to quantify attention allocation in mice in NOR-like paradigms. The findings obtained here should encourage future research to use odour-labelled in investigating the influence of emotion on attention and memory in tests.

## Supporting information

behavoiural data in experiments 1-7

## Acknowledgements

We thank G. Winocur for an inspiring discussion, J. Neill for her support, and C. Charalambous for statistical advice.

